# Targeting anemia-induced CD71^+^ reticulocytes protects mice from *Plasmodium* infection

**DOI:** 10.1101/2025.01.13.632761

**Authors:** Sareh Zeydabadinejad, Jong Sung Anthony Kim, Anna Zheng, Mrunmayee Rajendra Kandalgaonkar, Prince Boakye Ababio, Matam Vijay-Kumar, Beng San Yeoh, Piu Saha

## Abstract

Malaria, caused by *Plasmodium* spp., is a global health concern linked to anemia and increased mortality. Compensatory erythropoiesis seen during acute anemia results in an increased circulating reticulocyte count (*i.e.*, immature RBC) a key factor in understanding the relationship between pre-existing anemia and *Plasmodium* burden. Reticulocytes in mice are marked by transferrin receptor (CD71^+^) and glycophorin A-associated protein (Ter119^+^). To model acute anemia with increased reticulocytes, C57BL/6 mice were either bled (*i.e.* phlebotomized) or administered phenylhydrazine, before being infected with *Plasmodium yoelii* (*P. yoelii*), a mouse-specific strain with a preference for reticulocytes. In *P. yoelii*-infected anemic mice, we observed heightened parasitemia and significant body weight loss compared with non-anemic *P. yoelii*-infected mice. Additionally, serum inflammatory cytokines, erythropoietin, and liver injury markers, along with hemozoin deposition significantly increased in anemic *P. yoelii*-infected mice. RBC transfusion from healthy normal donors to *P. yoelii*-infected anemic recipient mice ameliorated anemia by reducing overall reticulocyte count and increasing mature RBC count. RBC transfusion rescued body weight loss, decreased parasitemia, and reduced serum erythropoietin levels. Finally, to confirm the role of reticulocytes in *P. yoelii* infection, reticulocytes were depleted using anti-CD71 monoclonal antibody in *P. yoelii*-infected mice. We observed improvement in hematologic parameters and stark reduction in parasitemia in both pre-existing anemic and non-anemic *P. yoelii*-infected mice. Collectively, our results suggest that pre-existing anemia may increase the risk of *Plasmodium* infection due to the greater reticulocytes population. Anti-CD71 treatment in *Plasmodium* infection may offer a novel therapeutic strategy to combat *Plasmodium* infection and malaria.

**Grant support:** This work was supported by grants from the Crohn’s and Colitis Foundation (CCF) and American Heart Association (AHA) Career Development Award (854385 and 855256 respectively) to Piu Saha; grant from the National Institutes of Health (NIH) to Matam Vijay-Kumar (DK134053) and Liver Scholar Award from American liver Foundation to Beng San Yeoh.

## Introduction

Malaria is a vector-born disease caused by the protozoan parasite, *Plasmodium*. Despite the increasing efforts to minimize its public health burden in malaria-endemic regions, *Plasmodium* continues to infect over 245 million people each year globally and claimed 619,000 lives in 2021. *Plasmodium falciparum* and *Plasmodium vivax* are of the greatest concern, accounting for 95% of human malaria cases. *P. falciparum* is responsible for most malaria mortality, while *P. vivax* is the most frequent cause of malaria. The disparity in disease severity caused by the two strains is, in part, due to differences in their inclinations to infect immature RBC (reticulocytes) *versus* mature RBC (*aka* erythrocytes). Whereas *P. vivax* could only infect reticulocytes (1), *P. falciparum* infects both reticulocytes and erythrocytes (2). Despite the less restrictive host tropism of *P. falciparum*, the parasite exhibits a strong preference to infect reticulocytes when available (2). Thus, having a better understanding of the RBC tropism of *Plasmodium* spp. would help in the development of anti-malarial therapeutics tailored specifically to prevent RBC parasitization.

Reticulocytes are distinctively characterized by their surface co-expression of the glycophorin A-associated protein, Ter119 (*i.e.*, erythroid lineage marker) and the transferrin receptor, CD71, that plays a key role in iron uptake (2-4). As the reticulocytes reach terminal maturation, they not only remove their nucleus and organelles, but also shed CD71 from their cell membrane (5). Circulating reticulocytes which retain CD71 represent only a small fraction of the total RBC pool, as most RBC matured in the bone marrow before they are released into the bloodstream (5). However, in the event of blood loss and/or anemia, the bone marrow is unable to adequately replenish the RBC pool, and this activates the compensatory stress erythropoiesis (*alias* extramedullary hematopoiesis) in the spleen and liver. The RBC produced in this manner are released prematurely, thus substantially elevating reticulocyte counts in circulation. Whether such an increase in circulating reticulocytes could predispose mice to severe malaria, particularly if it occurs before *Plasmodium* infection, remains poorly understood and warrants further study.

Severe anemia significantly contributes to malaria-associated pathology, particularly among children in sub-Saharan Africa which accounts for about 95% of global malaria cases, making it a major health concern (6-8). The onset of malarial anemia can be attributed to the lysis of parasitized RBC and the malaria-associated suppression of bone marrow erythropoiesis (*i.e.*, ineffective erythropoiesis) (8-10). Of note, most of the studies on malarial anemia are heavily focused on anemia being a consequence of the disease, rightfully so considering its role as a co-morbidity. In contrast, there are few studies that sought to determine whether pre-existing anemia could also play causal roles in increasing infectivity or susceptibility to *Plasmodium* infection. It is reasonable to consider, for instance, that individuals with higher reticulocyte count due to a pre-existing anemia may be more vulnerable to *Plasmodium* spp. that prefers to infect reticulocytes. Thus, in this study, we employed mouse models to investigate whether acute anemia that arises due to phlebotomy and/or phenylhydrazine (PHZ) could predispose mice to infection by *Plasmodium yoelii* (*i.e.*, a mouse-specific strain). Furthermore, we explored whether targeted depletion of reticulocytes via the use of anti-CD71 monoclonal antibody (α-CD71 mAb) could protect anemic mice from malaria.

Here we showed that phlebotomy and PHZ-induced anemia increased circulating reticulocytes and heightened the susceptibility to *P. yoelii* infection in mice. Malaria severity in pre-anemic mice was exacerbated as denoted by the increased parasitemia and markers of liver injury [serum total bile acids (TBA), alanine transaminase (ALT)], multi-organ injury [aspartate transaminase (AST)] and inflammation [lipocalin 2 (Lcn2), serum amyloid A (SAA)]. Transfusion of RBC into anemic mice infected with *P. yoelii* decreased the abundance of circulating reticulocytes and conferred protection against body weight loss, parasitemia, and other indices of malaria severity. Further depletion of CD71^+^ reticulocytes using α-CD71 mab in *P. yoelii*-infected mice, substantially improved the hematological parameters (RBC, hemoglobin and hematocrit), immune cell counts (WBC, monocytes, lymphocytes, neutrophils), and markedly reduced parasitemia in both pre-existing anemic and non-anemic mice. Collectively, our findings underscore the critical role of reticulocytes in malaria pathogenesis and propose novel therapeutic opportunities by targeting reticulocyte populations.

## Materials and methods

### Mice

C57BL/6J wild-type mice (Stock # 000664) were procured from Jackson Laboratory and bred in-house in the Department of Laboratory Animal Resources, University of Toledo College of Medicine and Life Sciences. Mice were maintained under specific-pathogen-free conditions, housed in cages containing corn-cob bedding (Bed-O-Cob, The Andersons Co.) and nestlets (Cat# CABFM00088, Ancare Corp.) and fed ad libitum grain-based chow (LabDiet 5001). Mice were housed at 23 °C with 12 h light/dark cycle. Age and sex matched mice (males and females) were used for all the experiments. Animal handling was conducted according to the Institutional Animal Care and Use Committee (IACUC) approved protocols.

### *Plasmodium yoelii* (*P. yoelii*) infection and quantification of parasitemia

The *P. yoelii* subsp. *yoelii*, strain 17XNL: PyGFP (Catalog No. MRA-817) (11), which stably expresses the green fluorescent protein (GFP), was acquired from BEI Resources and kept as frozen stocks of parasitized RBC. Mice were infected with *P. yoelii*-parasitized RBC delivering 1 x 10^5^ parasites in 100 µL sterile PBS through intraperitoneal (*i.p.*) injections on each side of the abdomen. Parasitemia was assessed via flow cytometry and was defined as the percentage of GFP-positive RBC in whole blood obtained from the tails of infected mice. The parasitemia in control and anemic mice was also determined by microscopy on Giemsa-stained thin blood films. At the termination of experiment, mice were euthanized via CO_2_ inhalation. The serum and organs were collected and stored at -80 °C for further analyses.

### Phlebotomy-induced anemia

Iatrogenic anemia was induced by submandibular phlebotomy of about 200 µL of blood from each mouse (n=5) on two consecutive days (12, 13). Mice were weighed prior to each blood collection. After confirming anemia via CBC, mice were infected with *P. yoelii*.

### Phenylhydrazine (PHZ)-induced hemolytic anemia

Phenylhydrazine hydrochloride (PHZ; Sigma, MO, USA) was prepared as 50 mg/mL stock solution in sterile PBS. Different concentrations of PHZ (10, 20 and 60 mg/kg bw) were diluted and administered via *i.p.* at two days prior to *P. yoelii* infection (14). The control group mice received the same amount of sterile PBS. For dose-dependent PHZ study, mice were injected either one or two doses of 60 mg/kg bw of PHZ. Mice in the control group received the same amount of PBS. The group receiving one dose of PHZ was infected with *P. yoelii* two days later. The group receiving two doses of PHZ was given the first dose 3 days before, and the second dose one day after, being infected with *P. yoelii*. Hematological analyses were performed to confirm anemia (*i.e.*, hemoglobin and hematocrit levels) prior to *P. yoelii* infection. Of note, PHZ induces the oxidation of hemoglobin, leading to its denaturation and precipitation, which results in the formation of fluorescent ‘*Heinz bodies’* within red blood cells. This fluorescence, detectable by flow cytometry, can confound the accurate measurement of GFP^+^ RBC (*P. yoelii* infected RBC), potentially leading to erroneous parasitemia readings (15, 16). To accurately estimate parasitemia in PHZ-treated mice, we subtract the readings (due to PHZ-associated RBC autofluorescence) of the uninfected subgroups from those of the infected groups. A dose-dependent increase in parasitemia was visually validated by preparing thin blood smears from both groups of mice and detecting GFP using FITC and Giemsa staining.

### Blood transfusion

Fresh whole blood from C57BL/6J wild-type donors (n=5; about 200 µL/mouse via submandibular phlebotomy) was collected in EDTA-coated microtubes. RBC pellet was collected and washed with sterile PBS. Approximately 10 × 10^8^ RBC were resuspended in 200 µL of sterile PBS and injected into *P. yoelii* infected mice (phlebotomy and PHZ-induced anemic) via tail vein on days 4, 6, and 8 *p.i*. Percent parasitemia and TER119^+^ CD71^+^ reticulocytes were assessed using flow cytometry.

### *In vivo* neutralization of CD71^+^ reticulocytes

Ten-week-old *P. yoelii*-infected WT female mice were treated with either anti-CD71 antibody (α-CD71 mAb; 200 μg/mouse, InVivoMAb, BioxCell) or isotype IgG control (BioxCell) at days 2, 4, and 6 *p.i.* Percent parasitemia and reticulocytes were determined using flow cytometry. A separate group of mice was subjected to phlebotomy-induced anemia (bled approximately 200 µL) and, after 24 hr, were administered the first dose of α-CD71 mAb (200 μg/mouse) or isotype IgG control. After another 24 hr, these mice were infected with *P. yoelii*. On days 2, 4, and 6 *p.i.*, mice were given the second, third, and fourth doses, respectively, of either α-CD71 mAb or Isotype IgG control. Percent parasitemia and reticulocytes were assessed via flow cytometry.

### Flow cytometric analysis of reticulocytes

Whole blood (∼2 µL) was collected via tail nick and suspended in 1 mL of sterile PBS (pH 7.0). Following centrifugation at 2000 rpm for 5 min, the supernatant was removed, and cells were resuspended in 1 mL PBS containing 0.2% FBS. After two additional washes and centrifugation, cells were resuspended in 100 µL PBS + 0.2% FBS. Subsequently, cells were stained with fluorophore-conjugated anti-mouse monoclonal antibodies (Ter119-PE, CD71-APC, CD71-FITC, BD Biosciences) in staining buffer (PBS + 0.2% FBS) and incubated for 40 min at room temperature in the dark. Following a final wash, the stained cells were analyzed using the Accuri C6 flow cytometer (BD Biosciences) and BD Accuri C6 Software.

### Serum collection

After euthanasia, blood was collected via heart puncture into BD microtainer tubes (BD Biosciences) and centrifuged at 10,000 rpm for 10 min. Hemolysis-free sera was collected and stored at -80 °C until further analyses.

### Hematological analyses

On day 0, prior to *P. yoelii* infection, blood was collected in EDTA-coated microtubes (Sarstedt Inc.) for a complete blood count (CBC) using an automated hematology analyzer VETSCAN HM5 Hematology Analyzer (ABAXIS HM5C & VS2, Allied Analytic). Parameters determined included WBC and RBC counts, hemoglobin concentration, hematocrit, mean corpuscular volume (MCV), mean corpuscular hemoglobin (MCH), and mean corpuscular hemoglobin concentration (MCHC).

### Serum biochemical analyses

Serum total bile acids (TBA) was measured using Diazyme total bile acids assay kit (Cat # DZ042A) as per manufacturer’s instructions. Furthermore, serum total cholesterol, aspartate transaminase (AST), Alanine aminotransferase (ALT) was measured using assay kits from Randox (Cat # CH200, AS3804, and AP9764, respectively) as per manufacturer’s instructions.

### Serum ELISA

Serum lipocalin 2 (LCN2, Cat # DY1857), serum amyloid A (SAA, Cat # DY2948-05), tumor necrosis factor alpha (TNF-α, Cat # DY410-05), interferon-gamma (IFN-γ, Cat # DY485), interleukin 10 (IL-10, Cat # DY417), interleukin 17 (IL-17, also known as IL-17A, Cat # DY5390), granulocyte colony-stimulating factor (G-CSF, Cat # DY414), keratinocyte-derived chemoattractant (KC, alias CXCL1, Cat # DY453), mouse erythropoietin (EPO, Cat # DY959) were measured in hemolysis-free serum via DuoSet ELISA kits from R&D Systems.

### Giemsa staining

Thin blood smears of uninfected and infected mice were fixed in methanol (Sigma-Aldrich) for 1-2 min and air dried. Next, the slides were submerged into Giemsa stain (1:20 in deionized water) (Sigma-Aldrich) for 15 min before they were rinsed in deionized water and air-dried for imaging and analysis using a Leica DM2500 LED optical microscope.

### Fluorescence microscopy

For fluorescence imaging, blood smears were fixed in methanol, imaged and analyzed using a BioTek Cytation 5 Cell Imaging Multimode Reader (Agilent microscope). All images were captured at 40X magnification (Bright field and FITC).

### Histology

Livers and spleen sections from uninfected and infected mice were fixed in 10% neutral buffered formalin (NBF), embedded in paraffin and sectioned (2 µm), and stained with Hematoxylin & Eosin (H&E). Histological images were generated from VS120 Virtual Slide Microscope (Olympus) and OlyVIA software.

### Statistical analysis

The results are presented as mean ± SEM. Significance between two groups was assessed using Student’s t-test (unpaired, two-tailed), where p<0.05 was deemed significant. For comparisons among means of three or more groups, one-way ANOVA with pairwise multiple comparisons was employed. All statistical analyses were conducted using GraphPad Prism 9.0 software (GraphPad Software, Inc, La Jolla, CA).

## Results

### Iatrogenic anemia increased Ter 119^+^ CD71^+^ reticulocytes

Frequent blood draws in a clinical setting can result in iatrogenic anemia (alias nosocomial anemia) (12, 13). To model such anemia in mice, we performed submandibular phlebotomy on C57BL/6J mice for two consecutive days. Complete blood count (CBC) analyses at 24 hr after the second blood draw confirmed that phlebotomized mice had a significant decrease in RBC count, hemoglobin, and hematocrit compared to control mice (Fig 1A–C). Phlebotomized mice also displayed a decrease in hemoglobin content as indicated by mean corpuscular hemoglobin (MCH; Fig 1E) and mean corpuscular hemoglobin concentration (MCHC; Fig 1F), but no difference in mean corpuscular volume (MCV; Fig 1D) compared to control mice. Total white blood cell (WBC) count was moderately reduced, whereas monocyte and neutrophil counts were significantly reduced in phlebotomized mice (Fig 1G-I). Lymphocyte count, however, was comparable between phlebotomized and control mice (Fig 1J). These results indicate that the mice in the phlebotomized group were anemic with compromised innate immune cells.

**Figure 1.**
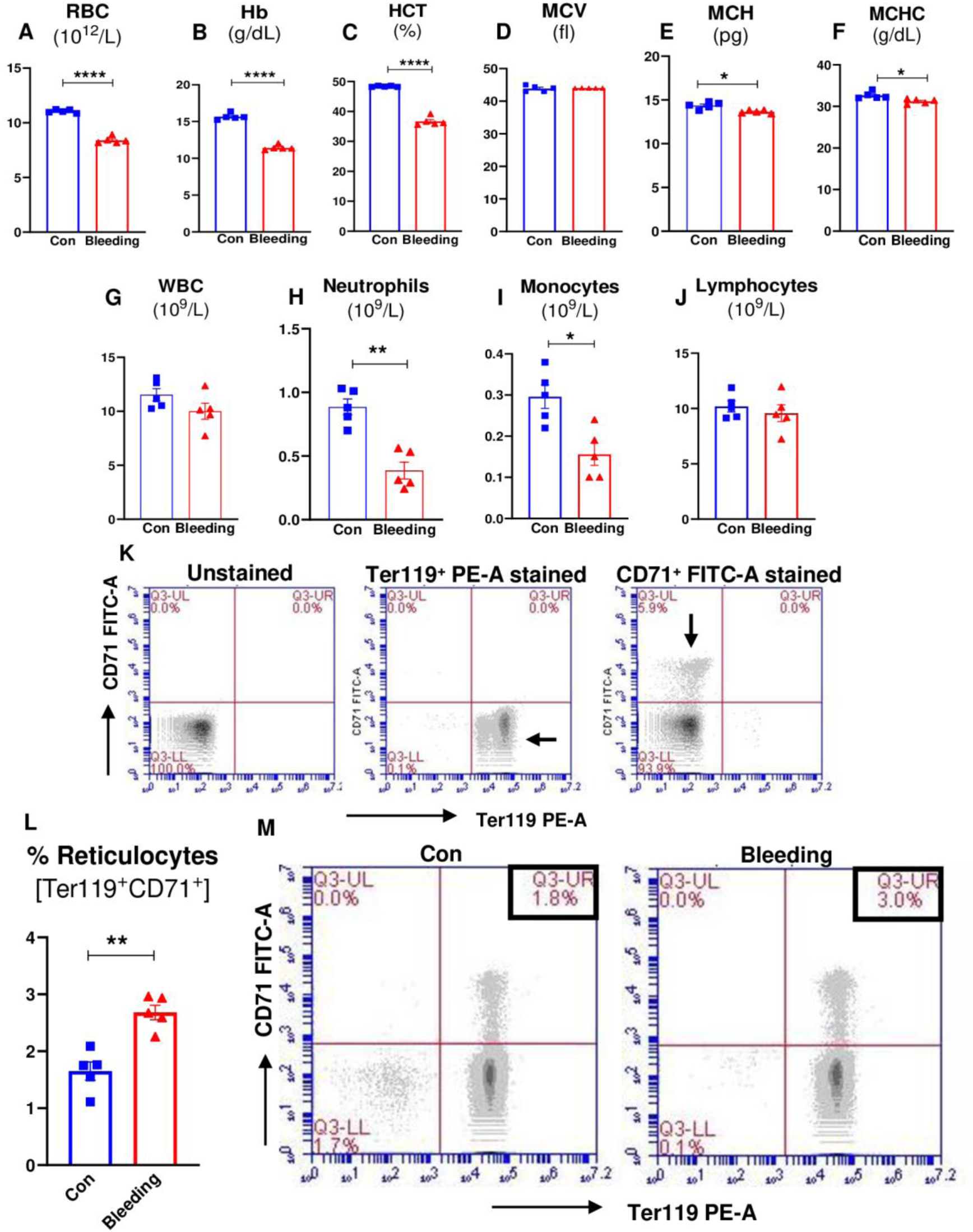
Iatrogenic (phlebotomy-induced) anemia lowered RBC, hematocrit and hemoglobin count and increased reticulocytes in mice. Blood (∼200 µL/ day) was collected from C57BL/6J wild-type (WT) mice (10-week-old males, n=5) via submandibular phlebotomy on two consecutive days to induce anemia. Blood from WT control and the phlebotomized mice (n=5/ group) was collected (EDTA tubes) for complete blood count (CBC; via VetScan hematology analyzer). Results for: **(A)** Red blood cells (RBC); **(B)** Hemoglobin (Hb); **(C)** Hematocrit (HCT, the volume percentage of RBC in blood); **(D)** Mean corpuscular volume (MCV, average size of RBC); **(E)** Mean corpuscular hemoglobin (MCH average amount of hemoglobin in RBC); **(F)** Mean corpuscular hemoglobin concentration (MCHC, average concentration of hemoglobin in a given volume of RBC) via CBC. **(G)** White blood cells (WBC), **(H)** Neutrophils, **(I)** Monocytes, **(J)** Lymphocytes. Flow cytometry analysis for reticulocytes (Ter119^+^ CD71^+^ cells) in blood. Representative **(K)** gating strategy for plots single stained Ter119^+^ and CD71^+^ RBC. **(L)** Bar graphs for % reticulocytes. **(M)** Representative dot plots for reticulocytes. Data represented as mean ± SEM from three independent experiments. *p<0.05, **p<0.01, ***p<0.0001.

Acute anemia triggers compensatory stress erythropoiesis to increase the production of RBC. However, the RBC produced in this manner are often released into the bloodstream before they are fully mature. These immature RBC (alias reticulocytes) can be identified via their surface expression of CD71 (2, 17). To confirm whether iatrogenic anemia induces reticulocytosis, we examined the percentage of Ter119^+^CD71^+^ RBC (*aka* reticulocytes) in phlebotomized and control mice via flow cytometry. Fig. 1K demonstrates the gating strategy for reticulocyte enumeration. As anticipated, phlebotomized mice exhibited approximately 1.7-fold greater abundance of reticulocytes compared to controls (Fig 1L-M) indicating phlebotomy-induced anemia leads to an increase in the percentage of circulating reticulocytes.

### Iatrogenic anemic mice were susceptible *to P. yoelii* infection

Anemia is a known sequela of malaria, but whether pre-existing anemia could drive the severity of disease is not completely understood. To demonstrate that anemia can exacerbate malaria, we infected both control and phlebotomized anemic mice with *P. yoelii*. Anemic mice showed a sharp decline in body weight post-infection (*p.i.*) that correlated with significant increases in parasitemia over the span of day 4-6 *p.i.* (Fig 2A, B). Non-anemic mice initially exhibited similar loss in body weight up to day 4, but recovered thereafter (Fig. 2A), associated with only a modest rise in parasitemia (Fig 2A-C). The anemic *P. yoelii-*infected mice also showed a decrease in liver and spleen weight (Fig 2D, E) compared to their non-anemic counterparts. Additionally, the serum isolated from the anemic parasitized mice had a stark yellowish appearance, which may be a consequence of hemolysis and/or malarial hepatitis and accumulation of bilirubin (Fig 2F). Furthermore, fluorescence microscopy analysis of blood smears collected on day 6 *p.i.* revealed that anemic mice exhibited a greater proportion of parasitized RBC (Fig 2G). Therefore, these results demonstrate that iatrogenic anemia can predispose mice to higher parasite burden and malaria-associated pathology.

**Figure 2.**
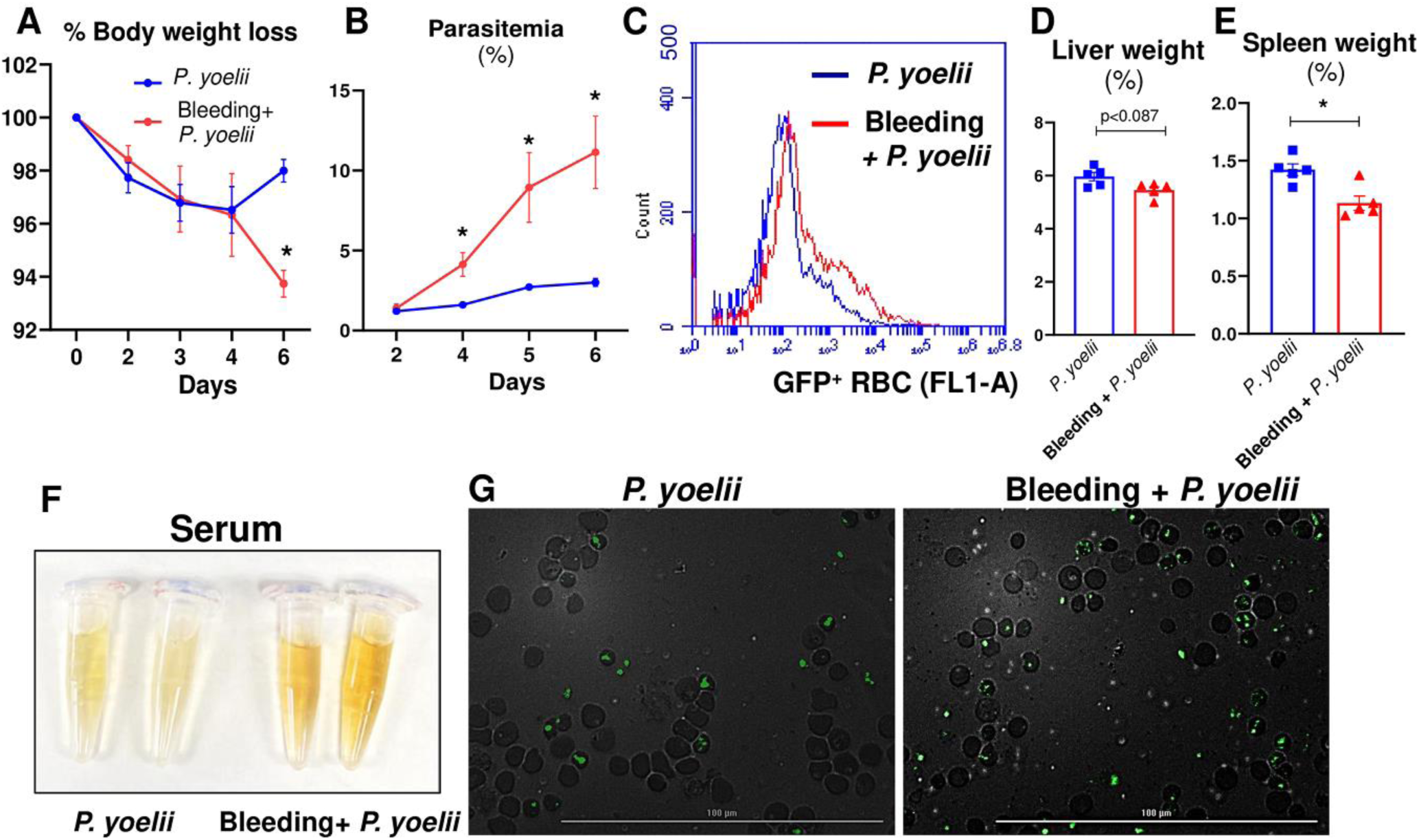
Iatrogenic anemic mice were susceptible *to P. yoelii* infection. Blood (∼200 µL/ day) was collected from WT (10-week-old males, n=5, two consecutive days) via submandibular phlebotomy to induce anemia. WT control mice and the phlebotomized mice (10-week-old males, n=5/ group) were infected with *P. yoelii*. (A) Body weight loss day 6 p.i. (B) % Parasitemia i.e. % GFP-positive RBC measured by flow cytometry during the infection. Quantification of parasitemia (% GFP^+^ RBC, i.e, GFP-*P. yoelii* infected RBC. (C) Flow cytometry representation of % GFP-*P. yoelii* infected RBC in histogram. (D) % Liver weight, (E) % Spleen weight, (F) Serum color 6 days p.i. (G) Representative images of GFP-*P. yoelii* infected RBC visualized under fluorescence microscope in *P. yoelii* infected mice 6 days p.i. Scale bar = 100 µm; magnification = 40X (Bright field and FITC). Data represented as mean ± SEM from three independent experiments. *p<0.05.

### *P. yoelii*-infected anemic mice suffered from severe hepatic injury and inflammation

The extent of liver damage has been shown to correlate with the severity of *Plasmodium* infection (18, 19). Accordingly, we sought to use the metrics of hepatitis markers to assess malaria severity among mice with and without iatrogenic anemia. While all *P. yoelii*-infected mice showed elevated levels of liver injury markers such as alanine transaminase (ALT) and total bile acids (TBA), these elevations were greater among mice with iatrogenic anemia (Fig 3A, B). Likewise, the level of the multiorgan injury marker AST was also significantly greater in anemic *P. yoelii*-infected mice (Fig 3C). Hematoxylin-eosin (H&E)-stained liver sections revealed substantial accumulation of hemozoin (a brown crystalline pigment produced by malaria parasites from the degradation of hemoglobin) (20) and prominent leukocyte infiltration in the liver of anemic *P. yoelii*-infected mice compared to their non-anemic counterparts (Fig 3D). Furthermore, spleen histology of the *P. yoelii*-infected anemic mice showed a considerable increase in splenic germinal centers relative to their non-anemic counterparts. *P. yoelii*-infected non-anemic mice maintained distinct regions of red pulp (RP) and white pulp (WP) in the spleen, while there was a marked loss of this distinction in their anemic counterparts (Fig 3E). Additionally, hemozoin in the spleen was more evident in anemic mice (Fig 3F). These histopathologic observations highlight significant differences in the liver and spleen in anemic versus non-anemic mice infected with *P. yoelii*.

**Figure 3.**
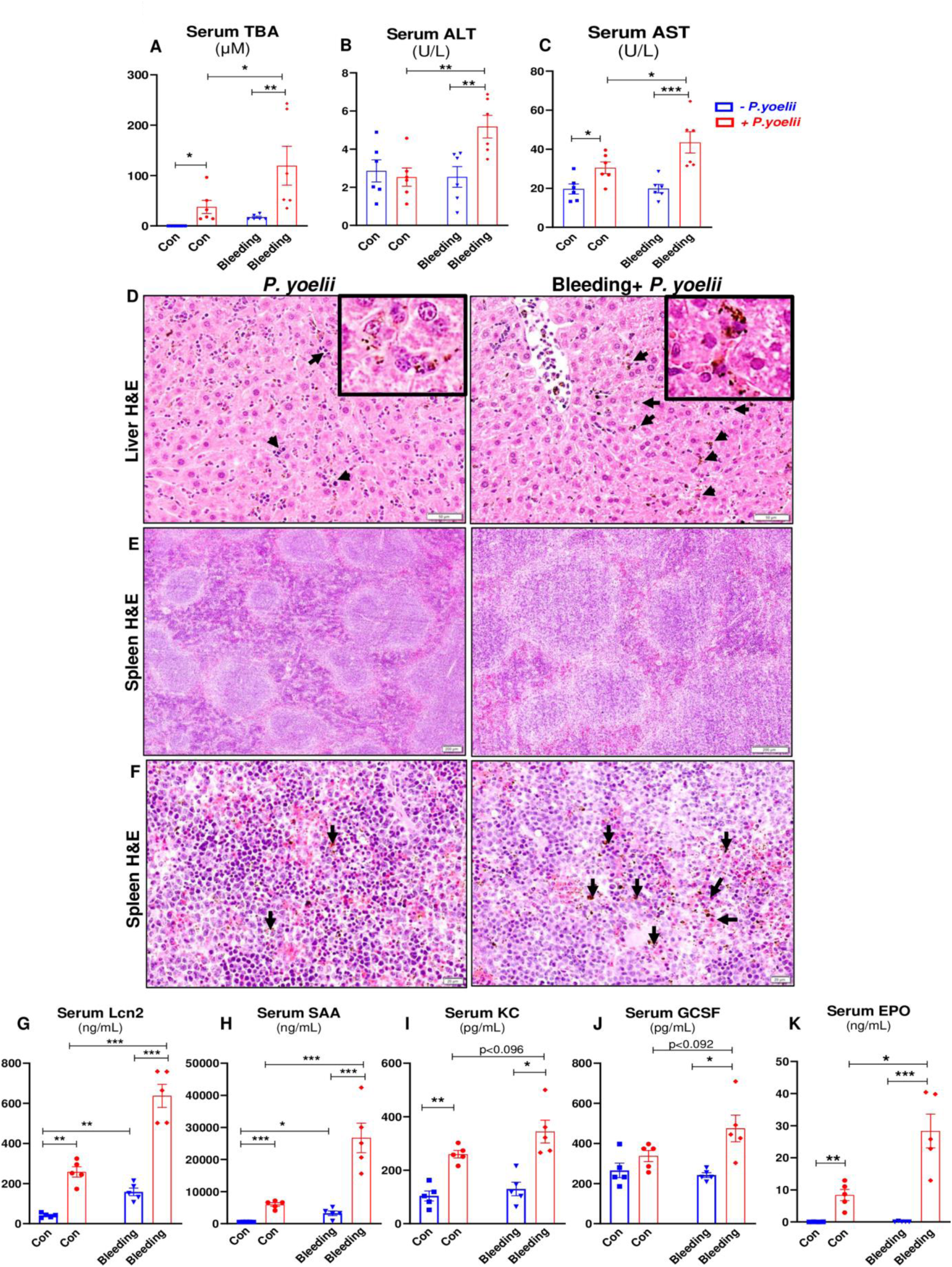
*P. yoelii* infected anemic mice displayed stark inflammatory responses. WT control mice and the phlebotomy-induced anemic mice (10-week-old males, n=6/ group) were infected with *P. yoelii*. Serum samples were collected on day 6 *p.i.* and analyzed for serum **(A)** total bile acids (TBA), **(B)** alanine transaminase (ALT), **(C)** aspartate aminotransferase (AST). Liver and spleen sections were processed for H&E staining for histopathological changes. **(D)** Liver histology Bars are 50 μm. *Inset:* parasitized red blood cells and hemozoin pigment bars are 20 μm. **(E-F)** Spleen histology: Bars are 200 μm and 20 μm. Black arrows indicated parasitized red blood cells (PRBC) and hemozoin pigment sequestration. Serum samples were collected on day 6 *p.i.* and analyzed for serum cytokines, **(G-J)** lipocalin 2 (Lcn2), serum amyloid A (SAA), KC (CXCL 1), granulocyte colony-stimulating factor (G-CSF) measured by ELISA. **(K)** Serum erythropoietin (EPO) measured by ELISA. Data represented as mean ± SEM from three independent experiments. *p<0.05, **p<0.01, ***p<0.001.

Next, we sought to assess the correlation between parasitemia and inflammation by measuring the inflammatory mediators in the sera collected on day 6 *p.i*. Both anemic and non-anemic mice infected with *P. yoelii* exhibited significantly elevated levels of the acute phase proteins lipocalin-2 (Lcn2) and serum amyloid A (SAA), as well as the chemokine keratinocyte chemoattractant (KC or CXCL1) (Fig 3G–I). Notably, Lcn2 and SAA levels were most prominently increased in anemic *P. yoelii*-infected mice (Fig 3G, H). The level of granulocyte colony stimulating factor (G-CSF, a growth factor promoting granulopoiesis) was moderately increased in the anemic *P. yoelii*-infected mice (Fig 3J). On the other hand, while serum erythropoietin (EPO; the hormone that stimulates erythropoiesis) was elevated significantly *p.i.* in both groups, the increase was modest in the non-anemic mice but striking in the anemic mice (Fig 3K). The greater increase in the latter may reflect a compensatory increase in RBC production in response to the anemic effect exerted additively by both phlebotomy and *P. yoelii* infection. Taken together, these results affirmed that iatrogenic anemia in mice can exacerbate malaria severity.

### Hemolytic anemia increased reticulocytes in mice

Next, we asked whether our findings could be recapitulated in a different model of anemia induced via hemolytic destruction of RBC by the chemical agent phenylhydrazine (PHZ) (14). Mice were injected with either a single dose of PHZ at various concentrations (10, 20, and 60 mg/kg b.w.), double dose of PHZ at 60 mg/kg, or PBS only. Blood samples were collected 48 hr post-administration from mice that received a single dose and 24 hr for double dose. CBC results indicate that RBC, hematocrit, and MCV decreased proportionally with increasing concentration of PHZ (Fig 4A–C). Interestingly, mice treated with a double dose of 60 mg/kg b.w. PHZ exhibited a significant increase in WBC, whereas a single dose notably decreased these cell populations (Fig 4D). Circulating neutrophils and monocytes increased with both single and double doses of 60 mg/kg b.w. PHZ (Fig 4E, F). Furthermore, a single dose of 60 mg/kg b.w. PHZ reduced lymphocytes, while double dose increased these cells (Fig 4G).

**Figure 4.**
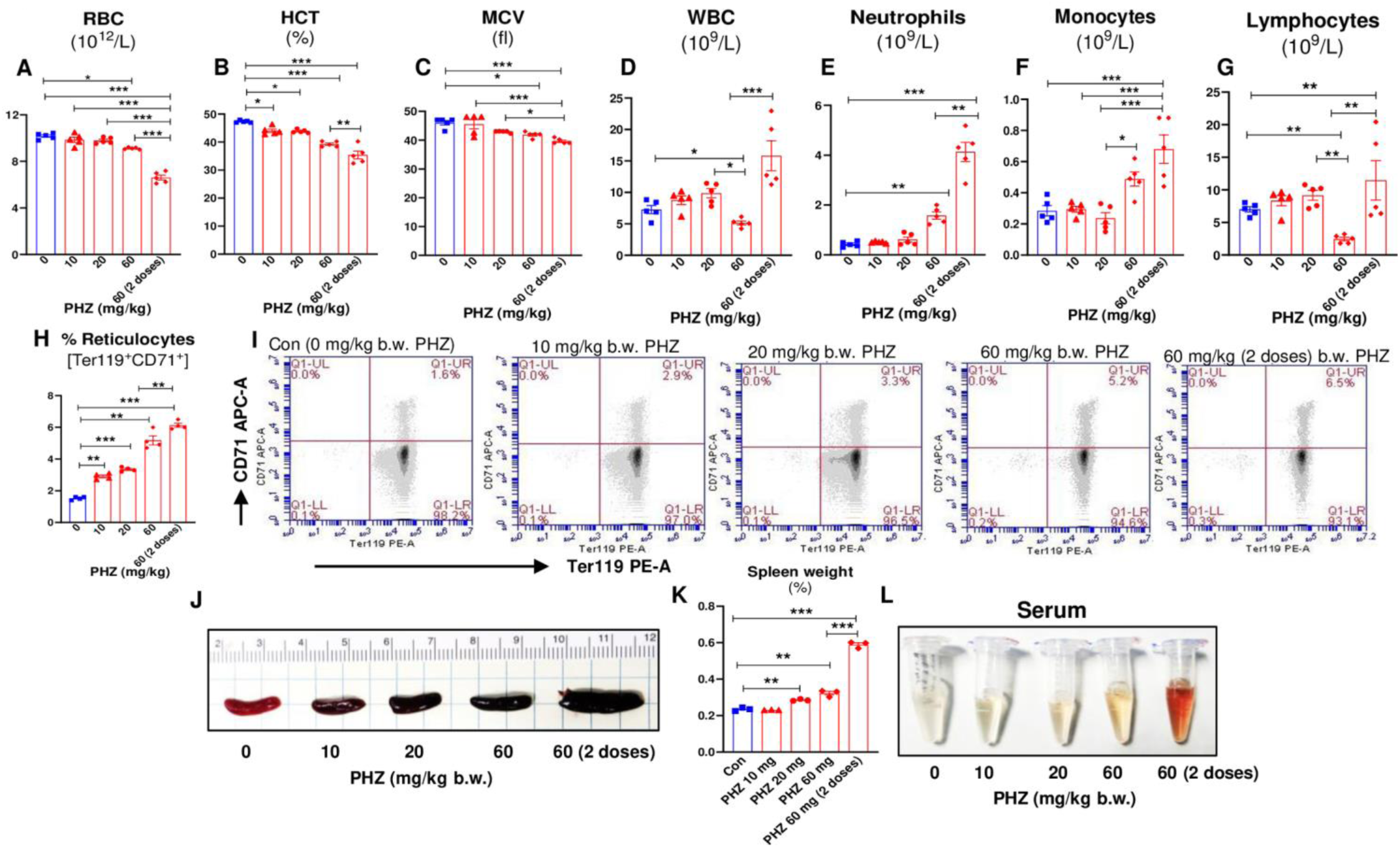
Phenylhydrazine (PHZ) induced hemolytic anemia in mice. PHZ [0, 10, 20 (1 dose) and 60 mg/kg bw. (1 and 2 doses)] were administered intraperitoneally (*i.p.*) to WT mice (10 weeks old, females) and after 48 hr PHZ injection, blood from WT control and the PHZ treated mice (n=5/ group, 10 weeks old, females) was collected (EDTA tubes) for CBC. Results for: **(A)** RBC, **(B)** HCT, **(C)** MCV, **(D)** WBC, **(E)** Neutrophils, **(F)** Monocytes, **(G)** Lymphocytes. Flow cytometry analysis for reticulocytes (Ter119^+^ CD71^+^ cells) in blood from 10-week-old females, WT control and the PHZ treated mice (10, 20 and 60 mg/kg bw*., i.p.* single and double doses). **(H)** bar graphs for reticulocytes. **(I)** Representative dot plots for reticulocytes after 48 hr PHZ treatment (dose dependent). **(J, K)** Gross spleen pictures and % spleen weight, **(L)** Serum color after 48 hr PHZ treatment (dose dependent). Data represented as mean ± SEM from three independent experiments. *p<0.05, **p<0.01, ***p<0.001.

PHZ treatment has been previously shown to induce reticulocytosis that peaks after 2–4 days (13, 21, 22). To determine the dose-response effect of PHZ on reticulocytosis, we collected blood at 48 hr post-PHZ and assessed the abundance of circulating Ter119^+^CD71^+^ reticulocytes. A proportional increase in circulatory reticulocytes was observed at 0, 10, 20, 60 (single dose), and 60 (double dose) mg/kg b.w. of PHZ (Fig 4H-I). Post-mortem examination at 48 hr after PHZ injection revealed significant splenic blackening and enlargement in mice that received 20 or 60 mg/kg (single or double dose, Fig 4J-K). The sera of mice treated with a double dose of 60 mg/kg PHZ exhibited a reddish-brown discoloration, indicative of pronounced hemolysis, therefore excluded from subsequent experiments.

### PHZ-induced anemic mice were highly susceptible *to P. yoelii* infection

To evaluate the impact of PHZ-induced hemolytic anemia on *P. yoelii* infection, mice were administered varying doses of PHZ (10, 20, and 60 mg/kg b.w.) or PBS and divided into two subgroups: one infected with *P. yoelii* and the other receiving PBS only. A rapid decrease in body weights across all groups was observed starting on day 4 *p.i. P. yoelii* infected PHZ-induced anemic mice exhibited a significant body weight loss 6-7 days *p.i.* especially in the 20 and 60 mg/kg groups. In contrast, the mice received 0 and 10 mg/kg b.w. PHZ began to regain body weight after day 7 (Fig 5A). Importantly, percent survival of *P. yoelii* infected PHZ 60 mg/kg b.w. < PHZ 20 mg/kg b.w. < PHZ 10 mg/kg = control *P. yoelii* infected groups (Fig 5A, B). PHZ treated anemic mice showed an increase in parasitemia from day 3 *p.i.* compared to non-anemic *P. yoelii* infected mice (Fig 5C). Notably, a positive correlation was observed between PHZ concentration and parasitemia (5C-E). Post-mortem (7-8 days *p.i.*) organ weights show significant loss of liver, spleen and lung mass in mice received both 20 and 60 mg/kg PHZ (Fig 5F-H). Moreover, Giemsa staining and fluorescent microscopy revealed a dose-dependent correlation, with severity of anemia associated with more extensive infected RBC 8 days *p.i.* (Figure 5I-J).

**Figure 5.**
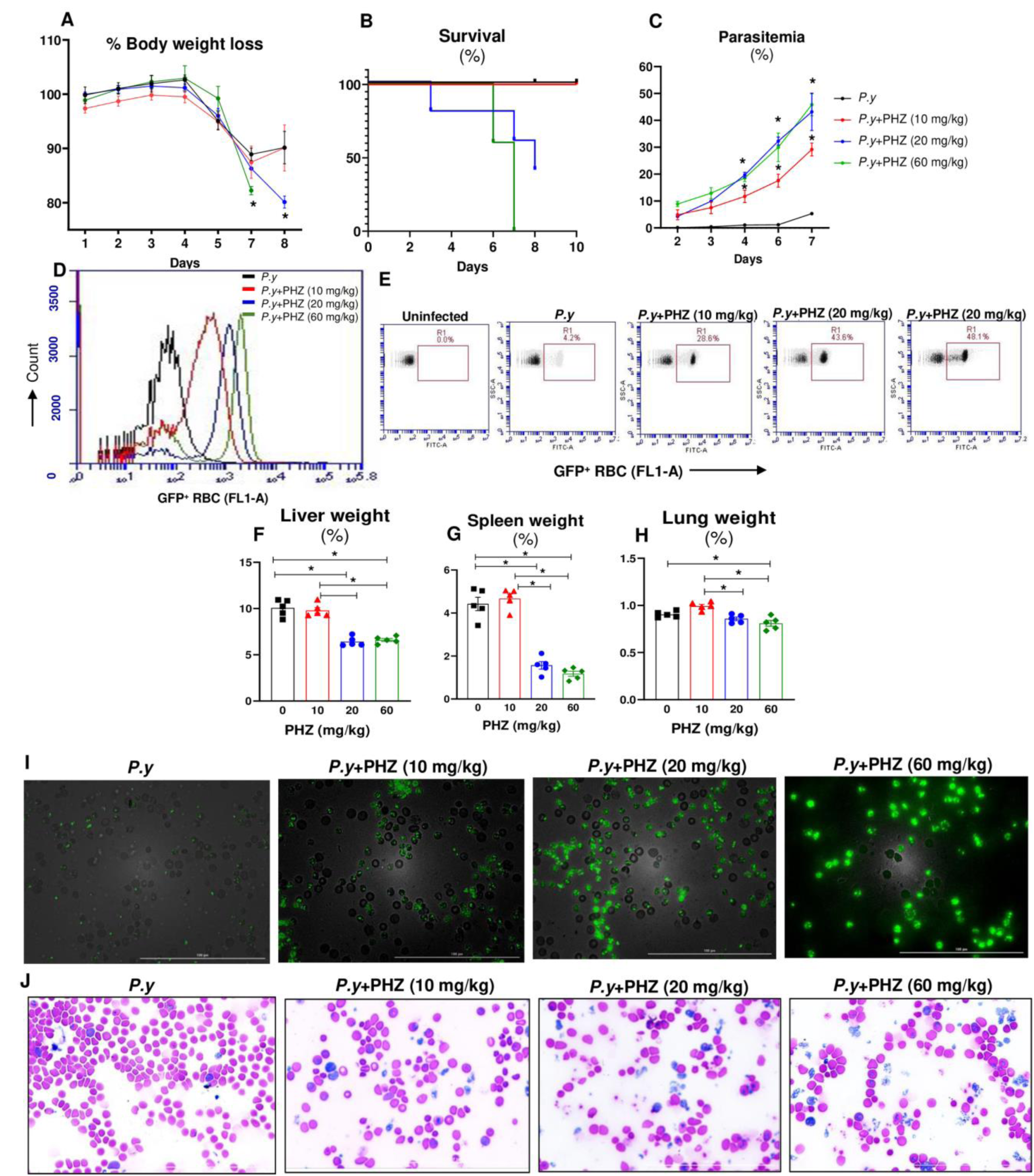
PHZ-induced anemic mice were highly susceptible *to P. yoelii* infection. WT control mice and the PHZ-induced (0, 10, 20 and 60 mg/kg bw., *i.p.* single dose, 48 hr after) anemic mice (10-week-old females, n=5/ group) were infected with *P. yoelii*. **(A)** % Body weight loss. **(B)** Survival graph. **(C)** % Parasitemia. **(D)** Flow cytometry representation of % GFP-*P. yoelii* infected RBC in histogram and **(E)** dot plots day 7 *p.i*., **(F)** % Liver weight, **(G)** % Spleen weight, **(H)** % Lung weight day 7-8 *p.i*., **(I)** Representative images of GFP-*P. yoelii* infected RBC visualized under fluorescence microscope in *P. yoelii* infected mice. **(J)** Representative images of infected RBC (Giemsa staining). Scale bar = 100 µm; magnification = 40X (Bright field and FITC). Data represented as mean ± SEM from three independent experiments. *p<0.05.

PHZ-induced anemic-*P. yoelii* infected mice exhibited heightened liver injury markers and elevated liver and spleen inflammation.

To assess liver injury in PHZ treated *P. yoelii*-infected mice, the principal serum markers of liver injury were measured. TBA, ALT and AST levels were significantly elevated in mice upon infection (Fig 6A-C) indicating severe hepatitis and multi organ damage in PHZ treated *P. yoelii*-infected mice. However, serum cholesterol levels were significantly reduced in all *P. yoelii* infected groups (Fig 6D) suggesting interrupted hepatic cholesterol biosynthesis (23). The results indicate that the degree of liver injury is positively associated with the degree of PHZ treatment and severity of infection.

**Figure 6.**
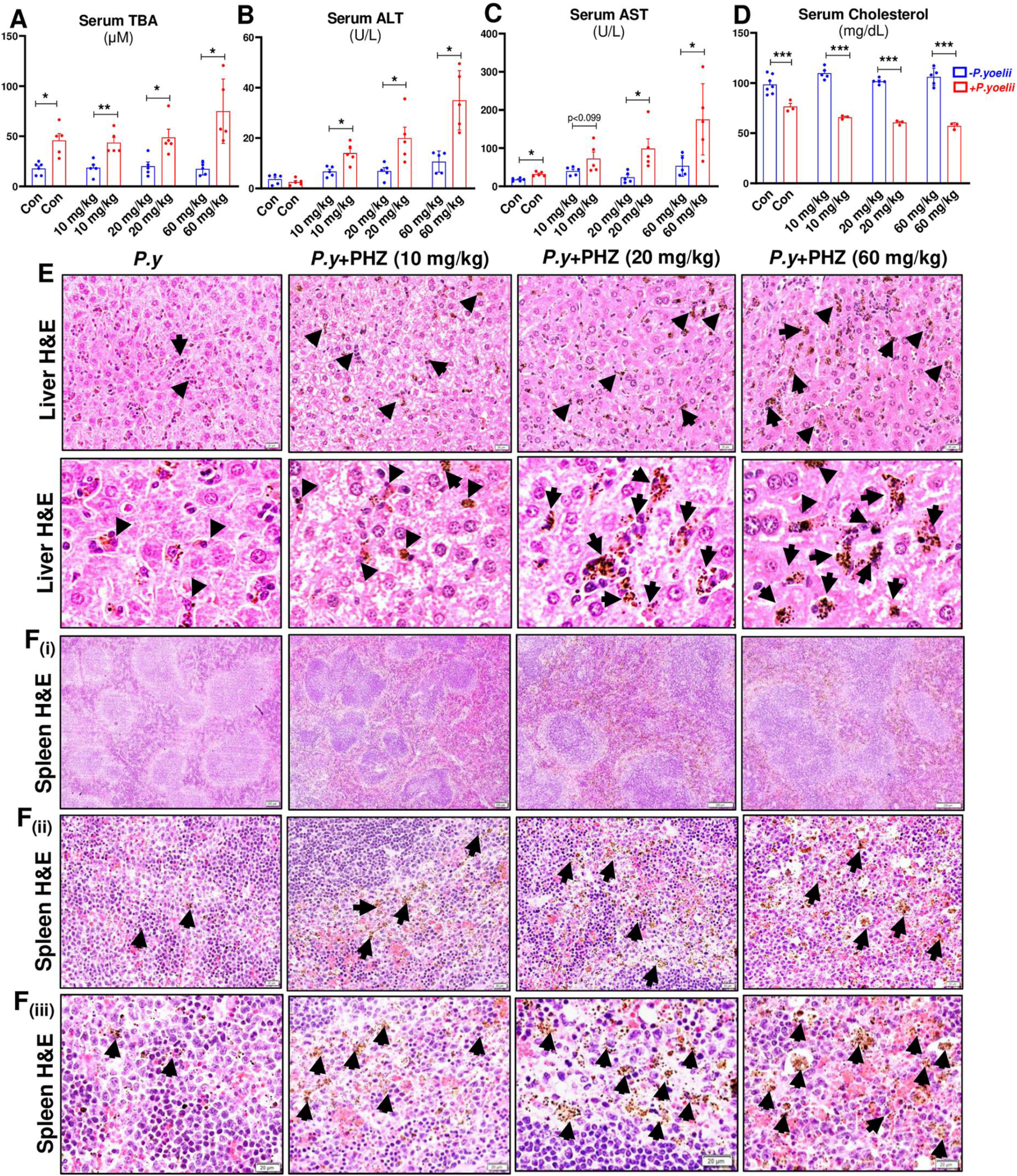
*P. yoelii* infected PHZ-induced anemic mice exhibited heightened liver and spleen inflammation. WT control mice and the PHZ-induced (0, 10, 20 and 60 mg/kg bw., *i.p.* single dose, 48 hr after) anemic mice (10-week-old females, n=5-6/ group) were infected with *P. yoelii*. Serum samples were collected 7-8 days *p.i.* and analyzed for Serum **(A)** TBA, **(B)** ALT, **(C)** AST and **(D)** cholesterol. Data represented as mean ± SEM from three independent experiments. 8 days *p.i.* the liver and spleen sections were processed for histopathological changes. **(E)** Liver histology: Bars are 20 μm. Spleen histology: Bars are **(F_i_)** 200 μm and **(F_ii_, F_iii_)** 20 μm. Parasitized red blood cells (PRBCs) and hemozoin pigments are marked with black arrows. *p<0.05, **p<0.01 and ***p<0.001.

Microscopic analysis of liver sections from days 7-8 *p.i.* revealed significant accumulation of hemozoin in PHZ-induced anemic mice. The degree of accumulation of hemozoin (black arrows) and immune cell infiltration in parasitized mice were PHZ 60 mg/kg b.w. ∼ PHZ 20 mg/kg b.w. > PHZ 10 mg/kg > control infection-only group (Fig 6E). Histological assessment of spleen on day 7 *p.i.* showed prominent loss of central germinal structure (Fig 6Fi) and hemozoin in PHZ (60 and 20 mg/kg) treated anemic mice compared with PHZ 10 mg/kg and control groups [Fig 6F (i, ii and iii)]. Histopathological analysis corroborated that *P. yoelii* infection induced more pronounced liver and splenic damage in mice with pre-existing anemia compared to non-anemic controls.

### PHZ-induced anemic *P. yoelii* infected mice showed stark increase in inflammatory responses

To determine the impact of the PHZ-induced anemia with *P. yoelii* infection on inflammation, serum levels of various proinflammatory cytokines and general inflammatory markers were assessed on both basal and the day of euthanasia (days 7-8 *p.i.*). Cytokines related to acute-phase responses (Lcn2 and SAA), neutrophil chemoattraction (KC), Th1-type responses (IFN-γ), regulatory immune responses (TNF-α), and erythropoiesis or hematopoiesis (IL-17, G-CSF and EPO) were measured. The results showed a significant elevation in Lcn2, SAA, and KC levels in all mice following *P. yoelii* infection, with a more pronounced increase in Lcn2 and SAA levels under anemic (Fig 7A-C) condition. Furthermore, there was a significant increase in serum levels of IFN-γ, TNF-α, and IL-17 in parasitized mice which received 60 mg/kg b.w. of PHZ (Fig 7D-F). In addition, G-CSF and EPO levels increased in mice that received either 20 or 60 mg/kg b.w. of PHZ upon infection (Fig. 7G, H). These results indicate that pre-existing hemolytic anemia aggravates the severity of inflammation during *P. yoelii* infection.

**Figure 7.**
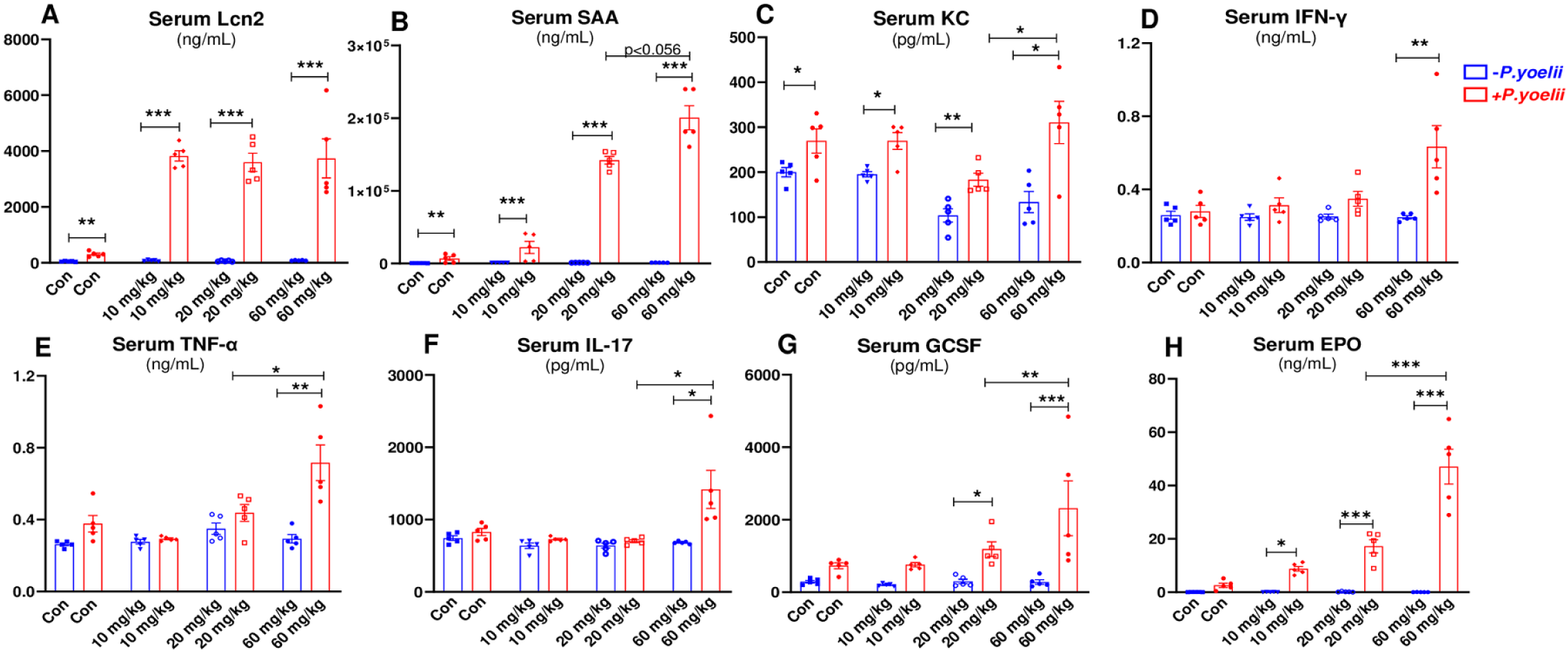
PHZ-induced anemic-*P. yoelii* infected mice showed heightened inflammatory cytokines. WT control mice and the PHZ-induced (0, 10, 20 and 60 mg/kg bw., *i.p*. single dose, 48 hr after) anemic mice (10-week-old females, n=5/ group) were infected with *P. yoelii*. Serum samples were collected 7-8 days *p.i.* and analyzed for serum cytokines, **(A)** Lcn2, **(B)** SAA, **(C)** KC, **(D)** IFN-γ, **(E)** TNF-α, **(F)** IL-17 and **(G)** G-CSF and **(H)** EPO measured by ELISA. Data represented as mean ± SEM from three independent experiments. *p<0.05, **p<0.01, ***p<0.001.

### RBC transfusion from normal donor mice ameliorated *P. yoelii* infection in pre-existing anemic mice by reducing the reticulocytes

Blood transfusion is often performed to treat severe anemia (24). Clinical studies on such intervention generally agree that its application can increase the survival rates of patients with severe malaria, presumably via the alleviation of anemia (24-27). How blood transfusion impacts *Plasmodium* infection itself, however, remains unclear. Given the profound impact of RBC transfusion in human malaria, we posited to perform a similar intervention in *P. yoelii*-infected mice with pre-existing anemia (induced via phlebotomy and PHZ treatment). We transfused RBC (10x10^8^ in 200 µL PBS, via tail vein) from healthy normal donor mice into *P. yoelii* infected anemic mice as illustrated in Fig 8A and supplemental Fig 1A. RBC transfusion protected *P. yoelii*-infected mice from further body weight deterioration (Fig 8B, supplemental Fig 1B) and reduced the percentage of parasitized RBCs (Fig 8C, supplemental Fig 1C). *RBC* Transfused mice began to recover after day 10 *p.i.* as indicated by their increasing body weights and decreasing parasitized RBCs (Fig 8B, C). Importantly, RBC transfusion significantly reduced reticulocyte counts in *P. yoelii*-infected anemic mice (Fig 8D, supplemental Fig 1D) which likely contributed to their protection from *P. yoelii* infection. Conversely, hematologic analyses confirmed a significant increase in RBC counts (Fig 8E, supplemental Fig 1E) and a marginal increase in hemoglobin (Fig 8F, supplemental Fig 1F) in RBC-transfused *P. yoelii*-infected anemic mice compared to *P. yoelii* infected anemic mice. There was also an increasing trend in hematocrit in RBC-transfused *P. yoelii* infected PHZ-induced anemic mice (supplemental Fig 1G), but no differences were observed in MCV and MCH values (supplemental Fig 1H, I). Furthermore, CBC results indicated a moderate reduction in the WBC and neutrophil counts, along with significantly reduced lymphocytes counts, in the RBC-transfused *P. yoelii* infected anemic mice (Fig 8G-I). Moreover, cholestatic and inflammatory markers were reduced in RBC-transfused *P. yoelii* infected groups compared to *P. yoelii*-infected anemic mice (Fig 8J-M and supplemental Fig 1K-L). Inflammatory markers, Lcn2 and SAA decreased notably in RBC-transfused *P. yoelii* infected anemic mice (Fig 8N, O). Additionally, serum EPO levels were significantly reduced in RBC transfused *P. yoelii*-infected mice, indicating that these mice were recovering from erythropoiesis (Fig 8P and supplemental Fig 1J). Histological analysis of liver and spleen sections showed a reduction in the accumulation of hemozoin and improved integrity of red pulp (RP) in RBC-transfused *P. yoelii* infected mice (Fig 8Q, R). These results suggested that RBC transfusion is a beneficial intervention in reducing parasitemia and ameliorating the effects of *P. yoelii* infection in anemic conditions, likely through mechanisms involving reticulocytes reduction and restoration of key hematologic parameters.

**Figure 8:**
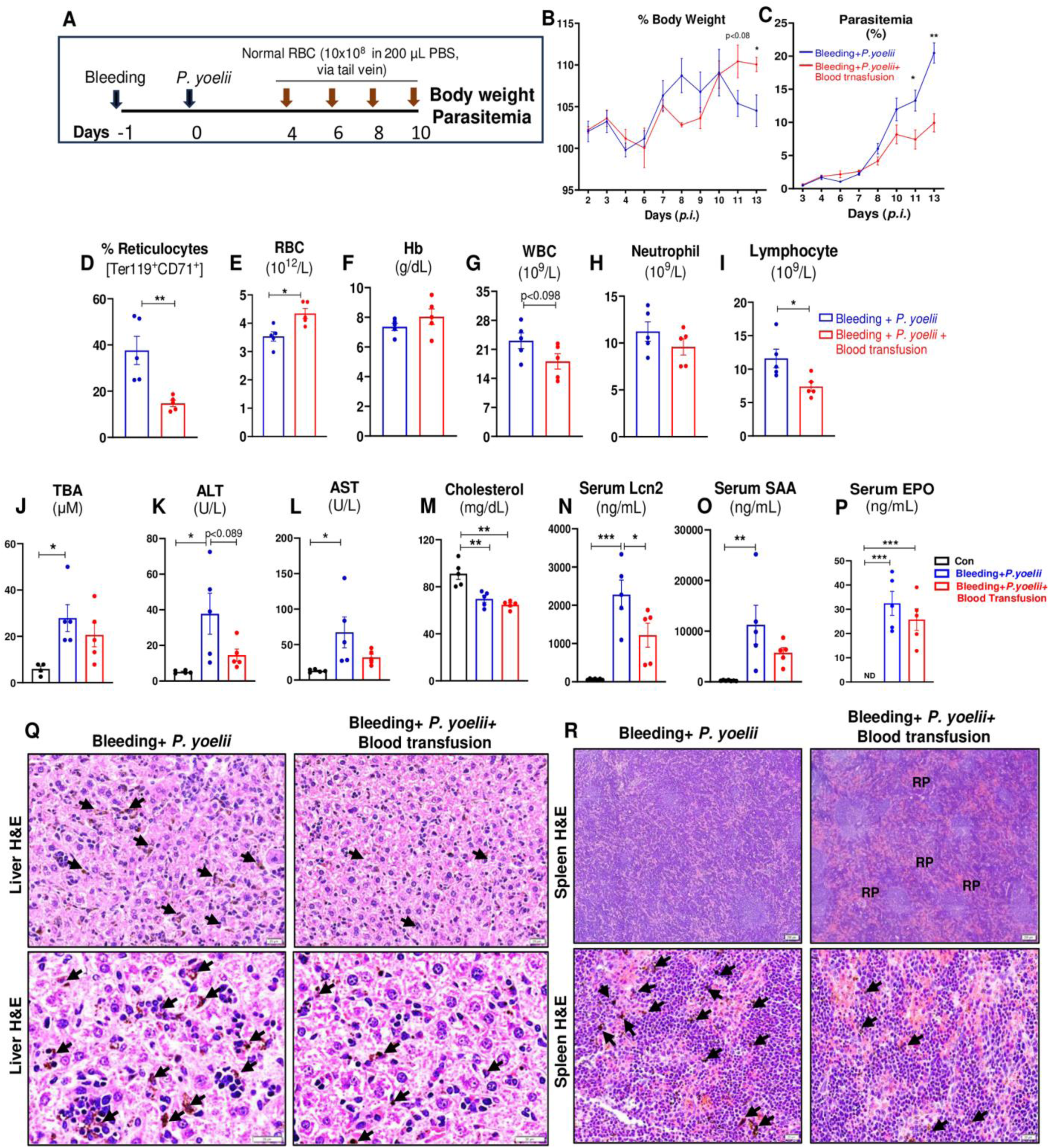
RBC transfusion reduced reticulocytes and ameliorated *P. yoelii* infection in phlebotomy-induced anemic mice. Phlebotomy-induced anemic mice (10-week-old females, n=5/ group) were infected with *P. yoelii*. Then, divided to two groups; one group received washed RBC (∼10 × 10^8^ RBC were resuspended in 200 µL of sterile PBS) from normal donors WT mice on days 4, 6, 8, and 10, next group received PBS *p.i*. **(A)** Experiment design. **(B)** % Body weight. **(C)** % Parasitemia *i.e.* GFP-positive RBC measured by flow cytometry during the infection. Quantification of parasitemia (% GFP^+^ RBC, *i.e*, GFP-*P. yoelii* infected RBC. **(D)** Reticulocytes (Ter119^+^ CD71^+^ cells) analysis day 13 *p.i.* Day 13 *p.i* CBC results for: **(E)** RBC, **(F)** Hb, **(G)** WBC, **(H)** Neutrophils, **(I)** Lymphocytes. Serum samples were collected 13 days *p.i.* and analyzed for serum **(J)** TBA, **(K)** ALT, **(L)** AST and **(M)** cholesterol. Serum cytokines, **(N)** Lcn2, **(O)** SAA and **(P)** EPO measured by ELISA. Data represented as mean ± SEM from three independent experiments. *p<0.05, **p<0.01, ***p<0.001.

### Depletion of CD71^+^ reticulocytes protected against *P. yoelii* infection

Reticulocytes are the primary targets for parasitization by many malarial parasites, including *P. vivax* and *P. yoelii*. Notably, *P. yoelii* demonstrates a 2.5-fold preference for reticulocytes over normocytes (2). One distinctive feature of reticulocytes is that they highly and selectively express CD71 (alias transferrin receptor 1) and would shed CD71 upon maturation (28). Hence, we sought to investigate whether anti-CD71 monoclonal antibody (α-CD71 mAb) could be leveraged to deplete circulating CD71^+^ reticulocytes and confer protection against *P. yoelii* infection. First, we determined that, upon *P. yoelii* infection, the percentage of circulating reticulocytes in mice significantly increased on day 2 and reached its peak at day 3-4 at ∼2-fold higher than baseline (Fig 9A). Thus, we opted to administer *P. yoelii*-infected mice with α-CD71 mAb starting on day 2, followed by repeated treatments on day 4 and 6 *p.i.* This intervention effectively depleted CD71^+^ reticulocytes, as confirmed by flow cytometry (Fig 9B). The liver and spleen of *P. yoelii*-infected mice treated with α-CD71 mAb displayed gross reduction in dark coloration compared to those treated with isotype antibody (Fig 9C). Although body weight differences were not initially observed, α-CD71 mAb treated *P. yoelii* infected mice began to regain weight on day 8 *p.i.* (Fig 9D). Notably, α-CD71 mAb treatment protected *P. yoelii*-infected mice from the progressive increase in parasitemia (Fig 9E). The anemia associated with *P. yoelii* infection, as indicated by the lowering of RBC counts, hemoglobin, and hematocrit *p.i.*, was also partially alleviated by α-CD71 mAb treatment (Fig 9F-H). Additionally, a substantial reduction in the WBC, neutrophil, monocyte and lymphocyte counts were observed, indicating decreased systemic inflammation in the α-CD71 mAb treated *P. yoelii* infected mice (Fig 9I-L). The inflammatory cytokines Lcn2, SAA and G-CSF decreased significantly in CD71^+^ reticulocyte-depleted *P. yoelii* infected mice (Fig 9M-O). Serum EPO levels also trended lower in α-CD71 mAb treated *P. yoelii-*infected mice (Fig 9P) in a manner reflecting the alleviation of malarial anemia (Fig 9F-H).

**Figure 9:**
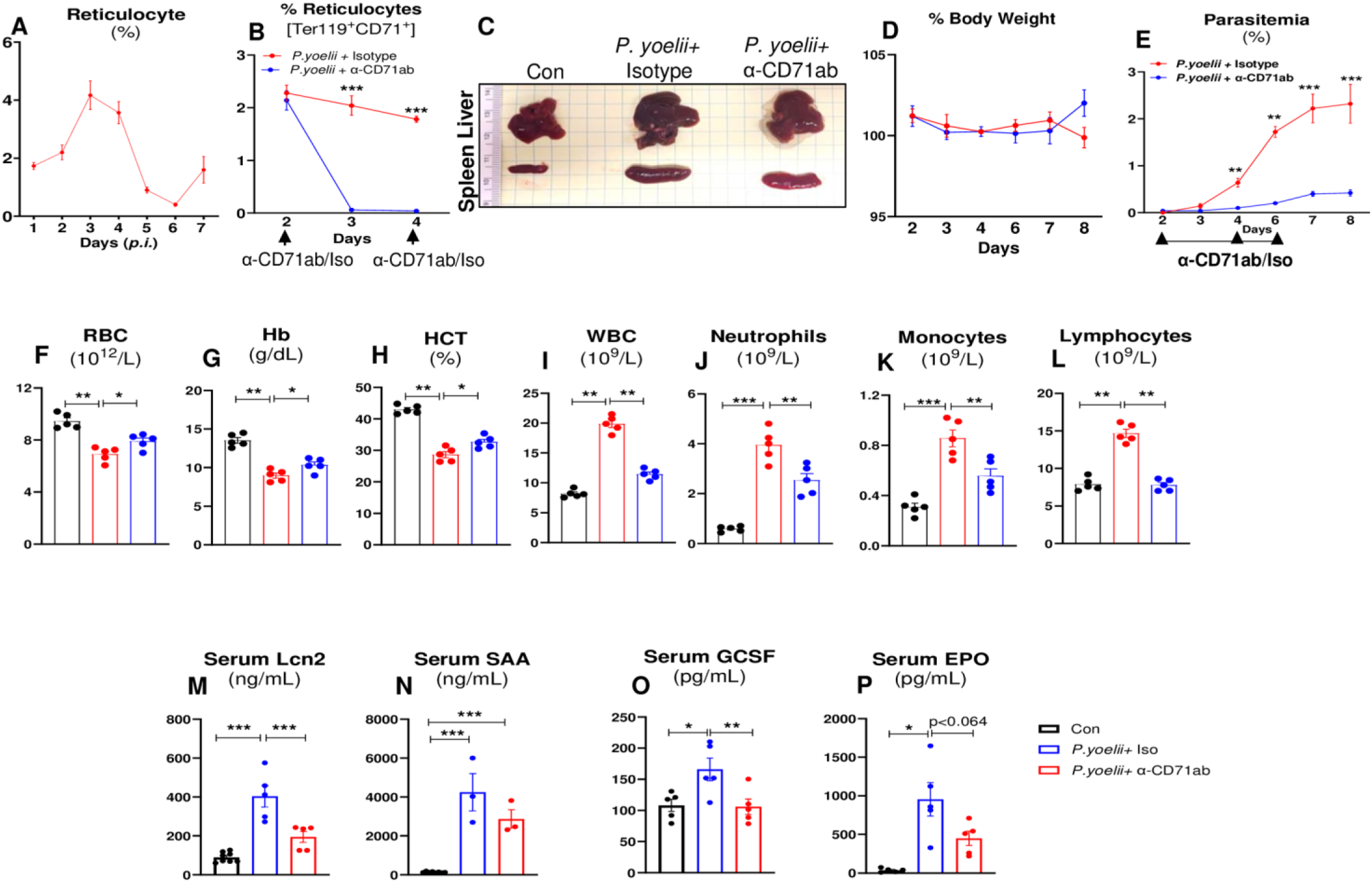
Depletion of CD71^+^ reticulocytes protected mice from *P. yoelii* infection. WT mice (10-weeks-old females, n=5/ group) were infected with *P. yoelii*. Then, divided to two groups; one group received anti-CD71 monoclonal antibody (α-CD71 mAb, TfR, 200 μg/mouse) on days 2, 4, and 6 and 2^nd^ group received isotype IgG control. **(A)** % Reticulocytes during progression of *P. yoelii* infection (days1-7 *p.i.*). **(B)** % Reticulocytes in *P. yoelii+* isotype and *P. yoelii*+ α-CD71 mAb treated groups. **(C)** Gross organ (liver and spleen) picture on day 8 *p.i*. **(D)** % Body weight. **(E)** % Parasitemia *i.e.* GFP-positive RBC measured by flow cytometry during the infection. Quantification of parasitemia (% GFP^+^ RBC, *i.e,* GFP-*P. yoelii* infected RBC. Blood from uninfected mice and *P. yoelii* infected mice that received either α-CD71 mAb or isotype was collected (EDTA tubes) on day 8 *p.i.* for CBC. Results for: **(F)** Red blood cells (RBC), Hemoglobin (Hb), **(H)** Hematocrit (HCT), **(I)** White blood cells (WBC), **(J)** Neutrophils **(K)** Monocytes, **(L)** Lymphocytes. Serum cytokines, **(M)** Lcn2, **(N)** SAA and **(O)** G-CSF and **(P)** EPO measured by ELISA 8 days *p.i.* Data represented as mean ± SEM from three independent experiments. *p<0.05, **p<0.01, ***p<0.001.

The decreased gross hepatic and splenic darkening in the α-CD71-treated *P. yoelii* infected mice compared to isotype-treated *P. yoelii*-infected mice (Fig 9C) suggesting that the protection was evident at the organ level as well. Our results indicated that α-CD71 mAb treatment significantly lowered both ALT and AST levels in *P. yoelii* infected mice (Fig 10A, B), affirming the alleviation of end organ damage *p.i.* Serum cholesterol levels, however, remained unchanged between the α-CD71 mAb treated and untreated groups (Fig 10C). Histological examination of liver sections on day 8 *p.i.* revealed a significant reduction in the accumulation of hemozoin and immune cell infiltration in α-CD71 mAb treated in *P. yoelii* infected mice (Fig 10D). Moreover, histological analysis of spleen showed a loss of central germinal structures, red pulp (RP) integrity, and hemozoin in isotype treated *P. yoelii* infected mice. In contrast, these pathological features were significantly diminished in α-CD71 mAb treated *P. yoelii* infected mice (Fig 10E). Collectively, our results underscore the significant impact of CD71^+^ reticulocyte depletion on the course of *P. yoelii* infection.

**Figure 10.**
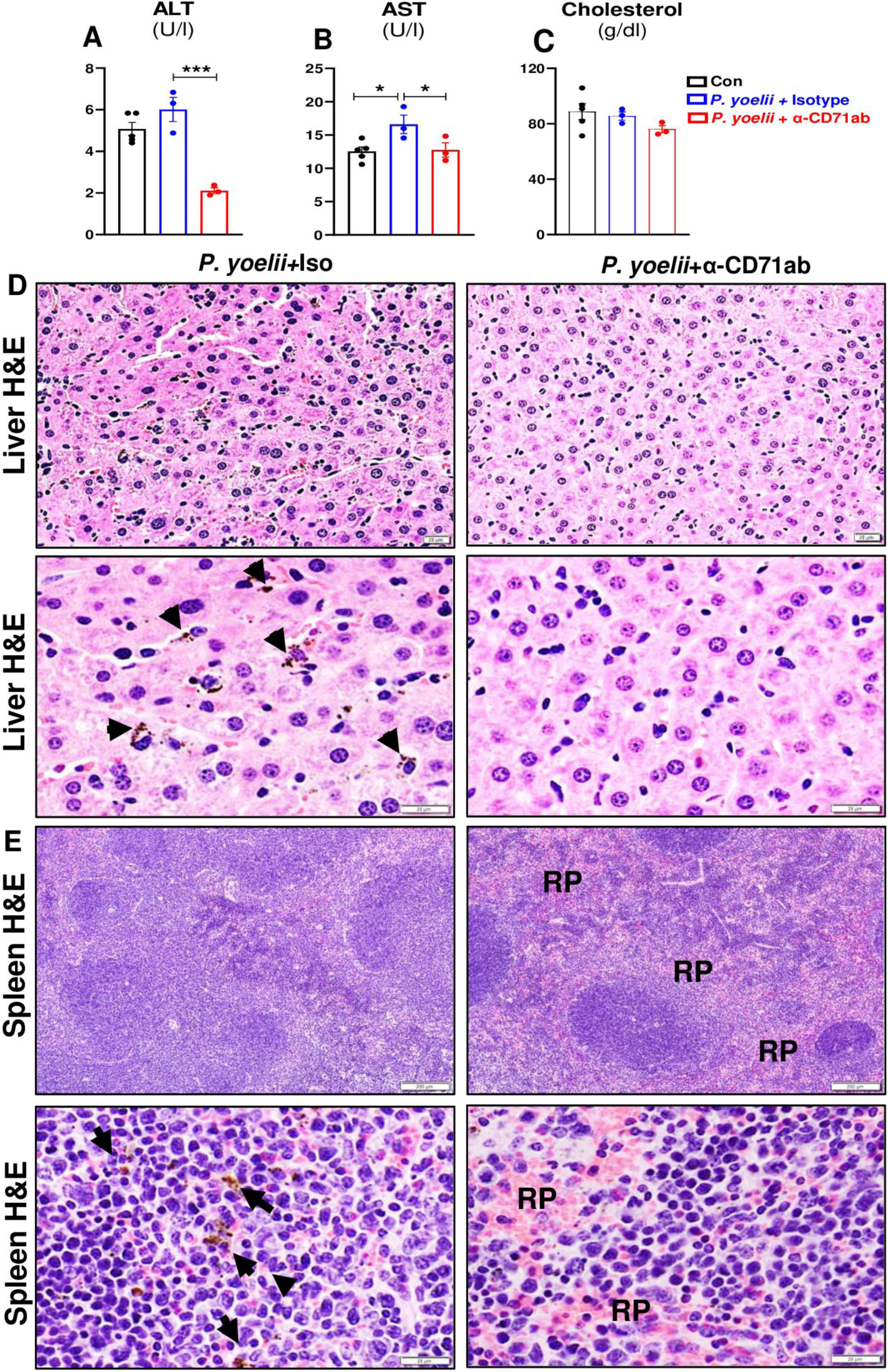
α-CD71 mAb treatment diminished liver and spleen inflammation in *P. yoelii* infected mice. WT mice (10-weeks-old females, n=5/ group) were infected with *P. yoelii*. Then, divided into two groups; one group received α-CD71 mAb (TfR, 200 μg/mouse) on days 2, 4, and 6 and 2nd group received isotype IgG Serum samples were collected 8 days *p.i.* and analyzed for serum **(A)** ALT, **(B)** AST and **(C)** cholesterol. 8 days *p.i.* the liver and spleen sections were processed for histopathological changes. **(D)** Liver histology: Bars are 20 μm, **(E)** Spleen histology: Bars are 200 μm and 20 μm. Parasitized red blood cells (PRBCs) and hemozoin pigments are marked with black arrows. RP = red pulp. Data represented as mean ± SEM from three independent experiments. *p<0.05, **p<0.01 and ***p<0.001.

Next, we aimed to determine whether targeted depletion of CD71^+^ reticulocytes could also protect mice that had pre-existing anemia before *P. yoelii* infection. To this end, we first induced anemia via phlebotomy, followed by α-CD71 mAb treatment and *P. yoelii* infection as illustrated in the experimental plan (Fig 11A). The subsequently observed body weights were comparable between α-CD71 mAb treated and isotype antibody-treated *P. yoelii* infected anemic mice (Fig 11B). Nonetheless, the administration of α-CD71 mAb significantly protected anemic mice from *P. yoelii* parasitemia compared with the isotype treated *P. yoelii* infected anemic mice (Fig 11C). In the isotype treated group, reticulocyte numbers initially increased on day 3 *p.i.* then decreased before increasing again after day 7 *p.i.* in a compensatory response to declining the RBC numbers (Fig 11D, blue line). Notably, despite the substantially diminished parasite load in the α-CD71 mAb treated mice, reticulocyte counts began to increase after 48 hr of α-CD71 mAb treatment, reflecting stress erythropoiesis due to depletion of reticulocytes (Fig 11D, red line). CBC results confirmed a significant increase in RBC, MCV and MCH values in the α-CD71 mAb treatment in anemic *P. yoelii* infected mice, while hemoglobin remained comparable in both treated and untreated groups suggesting recovery from anemia (Fig 11E-H). Additionally, notable reductions in the WBC, neutrophil, and lymphocyte counts were observed, indicating decreased systemic inflammation in the α-CD71 mAb treated *P. yoelii* infected anemic mice (Fig 11I-K). The systemic inflammatory markers Lcn2, SAA and G-CSF were also decreased significantly in α-CD71 mAb treatment in *P. yoelii* infected anemic mice (Fig 11L-N). Likewise, the serum EPO levels were significantly decreased in α-CD71 mAb treated *P. yoelii-*infected mice (Fig 11O), highlighting the protective effect of α-CD71 mAb treatment against *P. yoelii* infection in anemic condition. As was observed in the prior experiment, pre-anemic mice given α-CD71 mAb were protected from malaria-associated blackening of the liver and spleen, splenomegaly, and hepatomegaly (Fig 11P-R). Such protection was also evident at the histological level (Fig 11S-T). Taken together, these findings demonstrate the therapeutic potential of a targeted therapy against CD71^+^ reticulocyte, in the non-anemic and pre-anemic context, for the treatment of *P. yoelii* infection.

**Figure 11.**
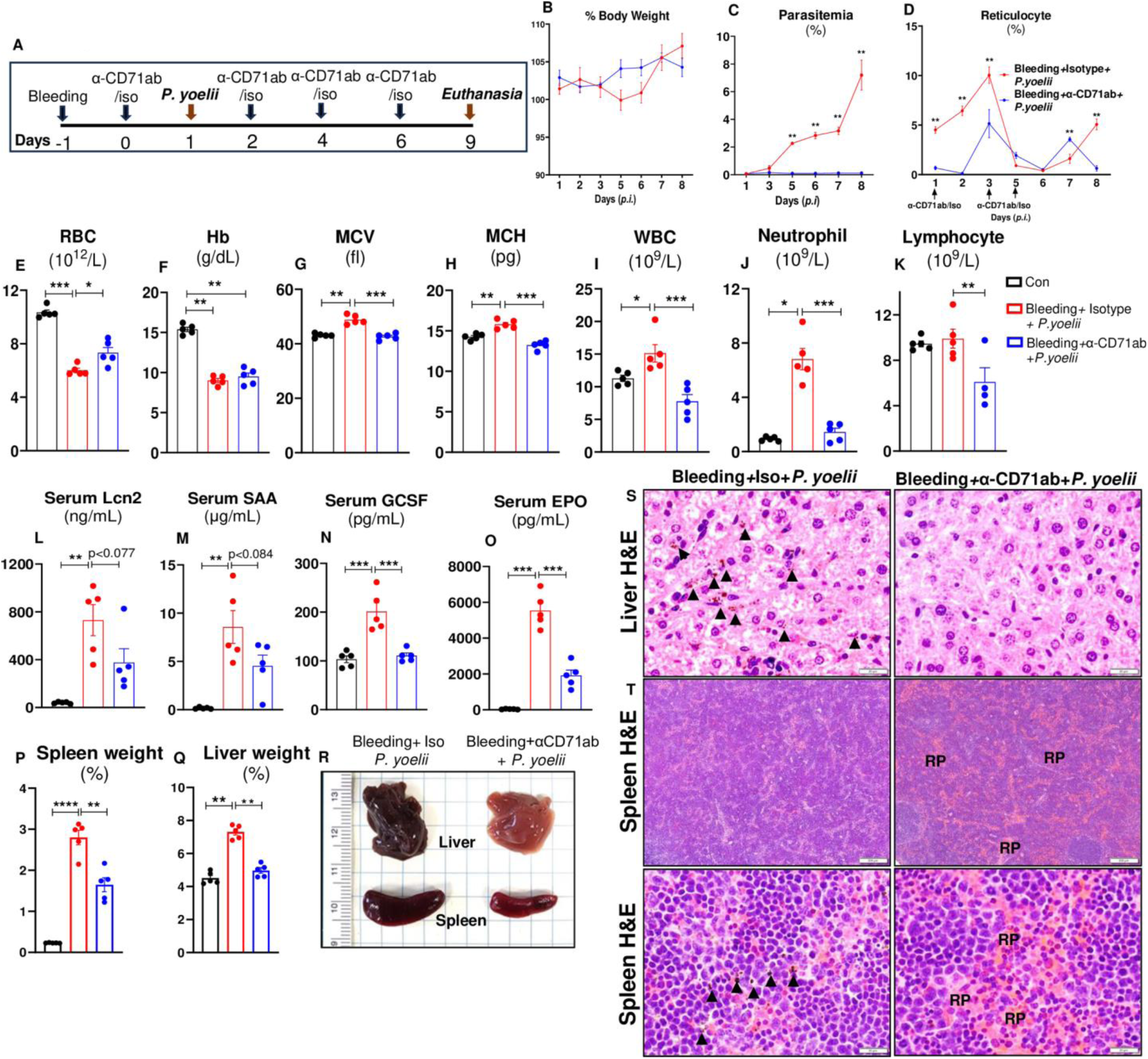
Depletion of CD71^+^ reticulocytes protected anemic mice from *P. yoelii* infection. Phlebotomy-induced anemic mice (10-week-old females, n=5/ group) received either α-CD71 mAb (TfR1) or isotype Ab on day 0, then infected with *P. yoelii* after 24h. **(A)** Overview of the study, **(B)** % body weight, **(C)** % Parasitemia *i.e.* GFP-*P. yoelii* infected RBC measured by flow cytometry during the infection. **(D)** % Reticulocytes (Ter119^+^ CD71^+^ cells). Blood from *P. yoelii* infected mice that received either α-CD71 mAb or isotype was collected (EDTA tubes) on day 8 *p.i.* for CBC. Results for: **(E)** RBC, **(F)** Hb (Hb), **(G)** MCV, **(H)** MCH, **(I)** WBC, **(J)** Neutrophils **(K)** Lymphocytes. Serum samples were collected on day -1 and 8 days *p.i.* and analyzed for serum cytokines, **(L)** Lcn2, **(M)** SAA, **(N)** G-CSF, **(O)** EPO. **(P)** % Spleen weight, **(Q)** % Liver weight. **(R)** Gross organ (liver and spleen) picture 8 days *p.i.* **(S)** Liver histology: Bars are 20 μm, **(T)** Spleen histology: Bars are 200 μm and 20 μm. Parasitized red blood cells (PRBCs) and hemozoin pigments are marked with black arrows. Data represented as mean ± SEM from three independent experiments. *p<0.05, **p<0.01 and ***p<0.001, ****p<0.0001.

## Discussion

Reticulocytes, or immature RBC, are the preferred target host cells for certain *Plasmodium* spp. that infect both humans and rodents (2, 17). For instance, *P. vivax* and the mouse-specific *P. yoelii* exhibit strong preferences for infecting reticulocytes (2). This selective tropism results from a complex interplay of host cell characteristics, parasite biology and immune evasion mechanisms. Reports have been demonstrated that reticulocytes provide several advantages to the parasites, including enhanced metabolic activity, increased protection from oxidative stress, and the ability to modify the reticulocyte membrane to better facilitate its intracellular expansion and immune evasion (2, 29). The circulating reticulocytes represent approximately 1-2% of the total RBC population in peripheral blood of a healthy individual. The tropism of *P. vivax* and *P. yoelii*, which are restricted to such a small RBC population in circulation, may explain, in part, their sub-lethal nature (2). On the other hand, *P. falciparum*, whose tropism is broader, are more likely to cause lethal infection (2). The importance of RBC tropism in determining malaria outcomes raises a prospective target that could curtail *Plasmodium* infection, thus alleviating malaria severity.

One prominent factor that can affect the proportion of circulating reticulocytes is anemia. Indeed, reticulocytes level is often used as a diagnostic marker for various types of anemia (2, 17). For instance, a low reticulocyte count is indicative of conditions such as pernicious anemia and iron deficiency anemia, whereas a high reticulocyte count suggests hemolytic anemia or anemia secondary to blood loss (2, 17). Taking into consideration the RBC tropism of *Plasmodium* spp., we reasoned that the anemia-associated increase in reticulocytes may aggravate *Plasmodium* infection. This may also be the case for the hypertensive BPH/2 mice, which, in a prior study, we found to display microcytic anemia, a greater reticulocyte population, and a more prolonged *P. yoelii* infection compared to the normotensive BPN/3 mice (30). To further determine the link between pre-anemic condition and malaria severity in this study, we employed phlebotomy and PHZ as tractable inducers of anemia in mice (13, 21, 31). Phlebotomy and PHZ treatment induced anemia and reticulocytosis in mice as anticipated, and more importantly, these anemic mice developed a more severe malarial pathology than control mice upon infection with *P. yoelii*. Consistent with our observation, the study by *Zhu et al*. found that PHZ-induced anemia can accelerate *P. berghei*-induced cerebral malaria in mice (21). Though our study employed different strains of mouse-specific *Plasmodium* spp., our findings concurred that pre-existing anemia could indeed pose as a risk factor for severe malaria.

Epidemiologic data directly linking iatrogenic or hemolytic anemia to malaria is currently lacking; however, such a link is apparent for the context of iron deficiency anemia. Intriguingly, iron deficiency anemia in humans is protective against malaria and, moreover, such protection could be negated by iron supplementation (32, 33). This is because, unlike hemolytic anemia, iron deficiency anemia presents defective reticulocytosis. It has been posited that the low reticulocyte count, rather than the anemia itself, is the primary factor that confers the resistance to malaria (32, 33). Iron supplementation rectifies such defective reticulocytosis, thus increasing reticulocyte population and reinstating the susceptibility to malaria (32, 33). While the type of anemia may differentially affect the predisposition to malaria, the underpinning link between reticulocytosis and malaria susceptibility is well-conserved and consistent across the types of anemia studied thus far (21, 32, 33).

Another line of evidence in support of the anemia-malaria link can be gleaned from the use of blood transfusion in clinical practice to manage severe malarial anemia (24-27). Clinical reports generally agree that blood transfusion is beneficial in improving the survival of patients with severe malaria. Previous studies on mouse models of malaria have also demonstrated that whole blood transfusion drastically increased survival rate and reduced parasitemia in mice infected with *P. chabaudi* and *P. berghei* (34-36). Likewise, our study indicates that RBC transfusion effectively mitigated symptoms of both phlebotomy-induced (Fig 8) and PHZ hemolytic anemia (supplementary figure 1) in *P. yoelii* -infected mice. Mice that received RBC transfusions exhibited accelerated weight gain, particularly following the third transfusion, and displayed reduced parasitemia. More importantly, RBC transfusion normalized the reticulocyte levels that were elevated by phlebotomy and PHZ-induced hemolysis. It is plausible that such protection could be attributed, in large part, to the suppression of reticulocytosis post-transfusion, which effectively deprives *P. yoelii* of reticulocytes to parasitize. Notwithstanding its benefit, blood transfusion is often performed in clinical settings with the intent to alleviate malarial anemia, rather than to treat *Plasmodium* infection *per se*. A change in paradigm with respect to the latter may benefit patient care, especially if further studies can demonstrate the efficacy of blood transfusion as a first line treatment rather than a delayed solution to malaria-induced anemia.

CD71 targeting therapy has been shown to be a promising anti-cancer treatment, since cancer cells overexpress CD71, *i.e.*, a transferrin receptor, to sustain their high iron demand (37, 38). Intriguingly, a recent study found that the anti-cancer efficacy of α-CD71 mAb could also be mediated via selective depletion of a pro-tumorigenic subset of CD71^+^ reticulocytes (39). Such a notion raises the possibility that α-CD71 mAb could be repurposed as a treatment for malaria, given that many *Plasmodium* spp. exhibit a strong tropism toward CD71^+^ reticulocytes. To our knowledge, this study is the first to provide the proof-of-concept on the use of α-CD71 mAb as an immunotherapeutic for treating *P. yoelii* infection. Of note, mice given α-CD71 mAb resisted the increase in parasitemia *p.i.* and were substantially protected from malaria-associated hepatitis. Mechanistically, α-CD71 mAb is likely to confer protection via depriving *P. yoelii* of its preferred host cell-type and/or by marking parasitized reticulocytes for destruction by immune cells. Intriguingly, the dose of α-CD71 mAb used herein was well-tolerated in mice and did not exacerbate anemia despite the consequent loss of reticulocytes. This is likely attributable to the short duration of the treatment and/or the depletion of primarily circulating reticulocytes, while sparing those undergoing maturation in key erythropoietic organs (*e.g.*, bone marrow). Further studies are warranted to address these hypotheses and to rule out the possibility of off-target or adverse effects. Further investigation into whether α-CD71 mAb can likewise curtail the infection caused by other species of *Plasmodium* would also be beneficial. Based on their varying tropism toward reticulocytes, it is reasonable to speculate that *P. chabaudi*, *P. berghei*, and *P. vivax* would be responsive to α-CD71 mAb treatment. Though the RBC tropism of *P. falciparum* is less restricted, its preference to invade reticulocyte would nonetheless make it susceptible to the effects of α-CD71 mAb. Thus, we envision that α-CD71 mAb has the potential to be harnessed, alongside other targets involved in RBC parasitization (*e.g.*, reticulocyte-binding 5 protein of *P. falciparum*) (40), as frontline medications for malaria.

Taken together, our study delineates the interplay among anemia, reticulocytosis and *P. yoelii* infection in mice. CD71^+^ reticulocytes were identified as the primary determinant of disease outcomes, whereby (i) its elevation due to anemia increases predisposition for severe malaria, and (ii) its suppression upon blood transfusion alleviates malaria severity. Most importantly, we demonstrate that antibody-mediated depletion of CD71^+^ reticulocytes confers profound protection against malaria. Future directions should include translating these findings into human malaria studies, where CD71^+^ reticulocytes also play a role in the disease. Investigating CD71-targeting approaches in human malaria could provide insights into novel therapeutic interventions, especially in cases of severe or life-threatening anemia.

## Acknowledgements

The authors would like to acknowledge technical support by Allen Schroering at the Histology Core of the Integrated Core Facility, the University of Toledo.

## Author Contributions

P.S. was responsible for conceptualization, formal analysis (with S.Z.), funding acquisition (with M.V.K.), project administration, resources (with M.V.K.), supervision, and visualization (with S.Z., J.S.A.K., A.Z., M.R.K., P.B.A., and B.S.Y.). Investigation was carried out by P.S., S.Z., B.S.Y., and M.V.K. Methodology was developed by P.S., S.Z., J.S.A.K., A.Z., M.R.K., P.B.A., and B.S.Y. The original draft was written by P.S., B.S.Y., S.Z., J.S.A.K., A.Z., and M.R.K. Review and editing were done by P.S., B.S.Y., S.Z., J.S.A.K., A.Z., M.R.K., P.B.A., and M.V.K.

