## Supplementary Fig1 for "Targeting anemia-induced CD71^+^ reticulocytes protects mice from *Plasmodium* infection"

**Supplementary Figure 1**

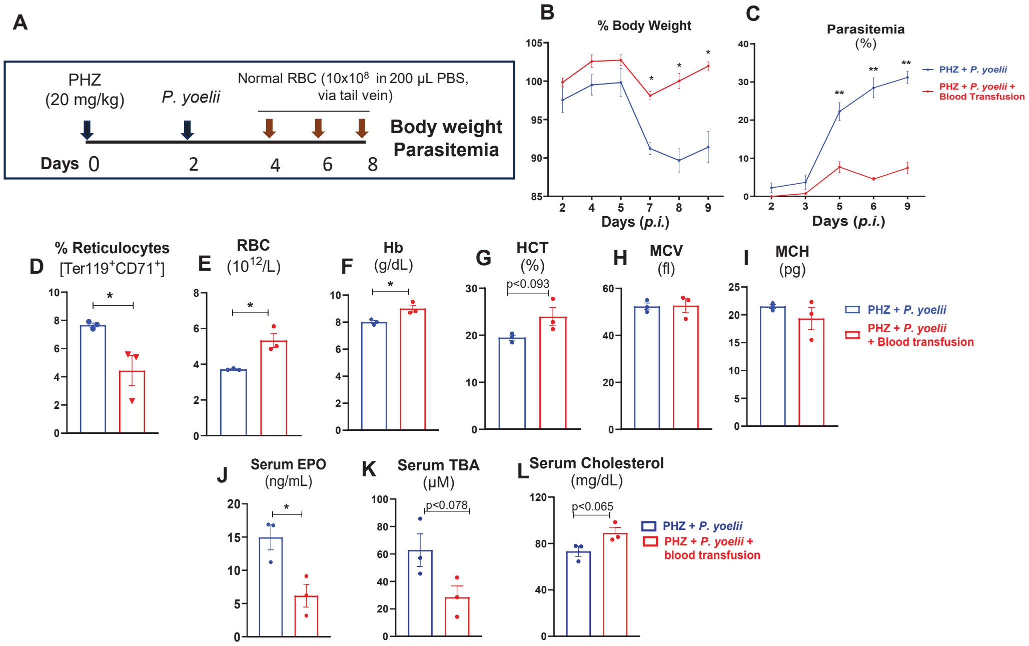

**Supplemental Figure 1: RBC transfusion decreased reticulocytes in circulation and mitigate *P. yoelii* infection in PHZ-induced anemic mice.**

PHZ-induced (20 mg/kg bw., *i.p.* single dose, 48h after) anemic mice (10-week-old males, n=5/ group) were infected with *P. yoelii*. Then, divided to two groups; one group received washed, one group received washed RBCs (~10 × $10^8$ RBC were resuspended in 200 µL of sterile PBS) from normal donors WT mice on days 4, 6, 8 next group received PBS *p.i.* **(A)** Experiment design. **(B)** % Body weight. **(C)** % Parasitemia *i.e.* GFP-positive RBC measured by flow cytometry during the infection. Quantification of parasitemia (% $GFP^+$ RBC, *i.e*, GFP-*P. yoelii* infected RBC. **(D)** Reticulocytes analysis day 9 *p.i.* Complete blood count (CBC; via VetScan hematology analyzer) day 9 *p.i.* Results for: **(E)** RBCs, **(F)** Hemoglobin (Hb), **(G)** HCT, **(H)** MCV and **(I)** MCH. Serum samples were collected 9 days *p.i.* and analyzed for serum **(J)** EPO, **(K)** total bile acids (TBA), **(L)** cholesterol. Data represented as mean ± SEM from three independent experiments. *p<0.05 and **p<0.01.
